# Water Hyacinth microbiome: metagenomic cues from environment and functionality in urban aquatic bodies

**DOI:** 10.1101/2023.03.09.531941

**Authors:** Rakeshkumar Yadav, Vinay Rajput, Mahesh Dharne

## Abstract

Water hyacinth (WH) is a widespread floating invasive aquatic plant having a prolific reproductive and dispersion rate. With the aid of its root-associated microbes, WH significantly modulates the ecosystem’s functioning. Despite their irrevocable importance, the WH microbiome remains unexplored in detail. Here, we present a shotgun analysis of WH rhizobiome (from urban rivers and a lake) and their surrounding water to unveil the diversity drivers and functional relationship. PCoA analysis revealed that microbial diversity of the WH is significantly shaped by the type of the aquatic bodies (River Vs Lake) (ANOSIM-R of 0.94 to 0.98 and R^2^ of 0.36 to 0.54). Temporal variations (River WH_2020 vs WH_2022) (R of 0.8 to 1 and R^2^ of 0.17 to 0.41) were observed in river WH, which could be mainly attributed to the transient taxa as there was higher sharing of core bacteria (48%). Also, the WH microbiome significantly differed (R= 0.46 to 1.0 and R^2^ of 0.18 to 0.52) from its surrounding water. WH inhabited more unique core members (42 to 45%), suggesting vertical transfer and selectivity in the microbiome. Functional metagenomics depicted the WH microbiome to harbour 140 pollutant-degrading enzymes involved in the degradation of various xenobiotic pollutants such as hydrocarbons, plastics, and dye. Moreover, an observed higher prevalence of metal and biocide resistance genes highlighted the persistence of resistant microbes assisting WH in environmental remediation application.

**Highlights:**

- Water hyacinth (WH) from urban water bodies inhabited by diverse microbial population
- First study to report variability in WH microbiome amid aquatic bodies and their environment
- Lake WH showed more unique core (43%), and River WH shared core bacteria (48%) across time
- 140 Pollutant degrading enzymes and 341 metal resistance genes indicates remediation potential

## 1. Introduction

Originated from the amazon basin of South America, the Water hyacinth (WH, *Eichhornia crassipes*, belonging to *Ponteneriaceae*) is a rapidly growing invasive floating macrophyte that has now been successfully introduced worldwide (Ávila et al., 2019; Xu et al., 2022; Villamagna and Murphy et al., 2010). The WH forms copious amounts of dense, interlocking mats on the surface of different aquatic bodies such as rivers, lakes and wetlands amid its prolific reproductive rate and root structure (Ávila et al., 2019; Xu et al., 2022). Due to their excessive growth, it completely covers the water bodies, preventing sunlight and oxygen from reaching the lower layers, thus making life disastrous for other aquatic flora and fauna (Galgali et al., 2023). Several strategies, such as mechanical removal and dredging, chemical control using herbicides, and biological control by pathogenic fungi and insects, have been employed to prevent the proliferation of WH (Xu et al., 2022). These efforts of removing WH are environmentally unsustainable and often need to be cost-effective.

Previous studies have mainly utilized WH in bioremediation as a phytoremediation-based removal of dyes, heavy metals, and other xenobiotic hydrocarbons owing to its high carbon, hydrogen and adsorbing qualities (Li et al., 2021; Delgado et al., 1993; Madikizela et al., 2021). Aquatic plant is a favourable ecological niche for microbial diversification and propagation, especially the roots that serve as hot spots for microbial diversity. These plants-associated microbes play a vital role in primary nutrient cycling, eliminating various pollutants, denitrification and phosphate uptake, influencing dissolved oxygen and generally have a mutualistic association with the host plants (Ávila et al., 2019; Achá et al., 2005, Pramanic et al., 2023). It is imperative to understand the WH-associated microbial population that significantly influences the overall nutrient cycling and biochemical process. Previous studies on the WH have provided insights into the role of these plant-associated microbes in assisting the removal of various pollutants and metals (Anudechakul et al., 2015; Achá et al., 2005). Further, a study by Luo et al., 2015, also reported a synergistic association of *Fusarium* with WH when exposed to heavy metals. Ávila et al., 2019, using 16S rRNA amplicon sequencing, gave a comprehensive understanding of the overall structure and functions of the WH rhizobiome in Amazon and Pantanal wetlands, highlighting the spatiotemporal changes. Another 16S rRNA-based study by Sharma et al., 2021, gave an account of the microbiome associated with WH of the Hindon River in India. These studies gave an overview of the WH microbiome and predicted putative functions based on the 16S rRNA study; however, the influence of the surrounding water and different aquatic bodies on the WH rhizobiome is not investigated so far. Although the role of WH is well-known in environmental bioremediation, detailed information on genes and enzymes for xenobiotic metabolism is lacking.

In the present study, we aimed to investigate the influence of surrounding water (environment) on the WH microbiome, how the WH microbiome differs from two different types of aquatic bodies (Lake vs River water hyacinths) undergoing different ecological processes, the temporal variations (changes in core and transient microbiome) in WH microbiome, and energy metabolism. Since WH is actively explored for environmental bioremediation, we also explored the potentials of the WH-associated microbiome to prove their role in assisting WH in remediation. To the best of our knowledge, this is the first shotgun metagenomic-based study to explain the relationship of WH microbiome with its surrounding environment, habitat, and its role in the bioremediation of toxic pollutants. Eventually, this study will help us to understand the propagation of WH in different aquatic bodies, especially in urban rivers and lakes, which shall have significant socio-economic impacts.

## 2. Materials and Methods

### 2.1. Sampling details

Pune city, one of the fastest-growing cities in the western part of India, is observing rapid urbanization and industrialization (Marathe et al., 2017; Yadav et al., 2021). Most rivers in Pune, such as River Mula, Mutha, Ramnadi, and Pawana, traverse the urban settlements and are in deteriorated conditions. These rivers are often plagued with luxuriously growing water hyacinth, thereby covering the entire surface and providing breeding grounds for mosquitoes resulting in a spike in dengue cases (Removal of water hyacinth, Citizens and water hyacinth). The local governing body in Pune recently announced more than 500 million INR for maintaining these invasive weeds (Removal of water hyacinth).

Here, we collected WH and corresponding samples from four sites (three river sites and one lake) from Pune city, India. The water hyacinth and the surrounding water samples were collected from three river sites (River Mula and Ramnadi) and one lake (Pashan Lake) from Pune city, India. The locations were named as A1 (Mula River, 18.565264, 73.788792), A2 (Ramnadi River, 18.548429, 73.79844), A3 (Mula River 18.567607, 73.796593) and WH (Pashan Lake, 18.535551, 73.787922). The sampling was performed in the month of March 2020, and the follow-up sampling was carried out in 2022 in the same month from the same sites. The average temperature in March in Pune city ranges from 30 to 35°C, and water hyacinth starts flourishing at these sites from January till the rainy season (June to August). In the first sampling, 12 WH samples were collected, triplicate from each site (A1, A2, A3, and WH), along with the four water samples (1 from each site). The WH was collected in sterile Hidispo bags (PW038, Himedia) and was immediately sealed and transported in dry ice. Around 1 litre of water was collected in sterile polypropylene bottles. In the second sampling, we collected 9 WH samples (triplicates from 3 sites, A1, A2, and A3) and three corresponding water samples. The sample details are stated in (Supplementary Table 1.). We did not conduct sampling from Pashan Lake in year 2022 due to inaccessibility.

### 2.2. Sample processing for DNA extraction

The water hyacinths were immediately transported to the laboratory for DNA extraction. The roots from all the plants were cleaned with sterile distilled water and cut into a uniform size of 2 cm. Further, these roots were gently washed in sterile distilled water twice to remove loosely attached microbial cells. The roots were crushed in the sterile mortar and pestle under sterile conditions. Around 250 mg of the roots of the DNA were processed for DNA extraction using DNeasy PowerSoil Pro Kit (Qiagen, 47014) as per the manufacturer’s instructions. The water samples were centrifuged at 7000 rpm for 5 minutes to remove coarse suspended particles. Further, the samples were subjected to successive filtration, first using a 0.4 uM pore size filter, followed by 0.22 uM filter paper (PES membrane filter, Himedia). The 0.22 uM filter was used for DNA extraction using the RNeasy Power water kit (Qiagen, 14700) as per the manufacturer’s instructions. The resultant DNA was analysed for purity (260/280) using NanoDrop (ThermoFischer NanoDrop Lite spectrophotometer), and concentration was estimated using Qubit dsDNA HS Assay Kit (Invitrogen, Q32851) using Qubit Fluorometer (Qubit 4 Fluorometer, Invitrogen).

### 2.3. MinION library construction and sequencing

The DNA with 260/280 purity of 1.7 to 1.9 was subjected to library preparation for nanopore sequencing. 1D Ligation sequencing kit SQK-LSK-109 was used for library preparation. Barcoding was carried out using EXP-NBD104 and NBD114. The library was prepared per the manufacturer’s instructions with the following modifications. The initial DNA concentration was 1.1ug/48ul for End-repair. The end-repair was performed at 20°C for 20 minutes, 65°C for 10 minutes and held at 4°C. The adapter and barcode ligation step was extended for 10-minute incubation than was mentioned in the procedure. Qubit was used to quantify the resultant library, and 450 ng was loaded onto FLO-MIN-106D, R9 Version.

### 2.4. Data analysis and statistics

The reads were basecalled and demultiplexed using Guppy software (V 5.0.11) (Wick et al., 2019). NanoFilt (v 2.7.1) was used for filtering and trimming of reads using a Q-score of 8 and read length of 500bp. The taxonomic classification was performed using the nr_euk database (released in Aril, 2020) using the Kaiju tool (Menzel et al., 2016) with default Greedy mode using an in-house customized script. Further, functions were analyzed using the KEGG database in MG-RAST (Keegan et al., 2016). The RemeDB (Sankara Subramanian et al., 2020) was used for analyzing pollutant degrading enzymes (PDEs) using the DIAMOND alignment tool (v 0.9.26) (Buchfink et al., 2015) using parameters for long reads (e-value of 10^−5^, identity of 60, --long reads (-F 15, -- range-culling, k 1), subject cover 40, --more-sensitive). Similarly, the BacMet database (v 2.0) (Pal et al., 2014) was used for biocide and metal resistance genes (MRGs) using the same diamond parameters. For taxonomic classification of genes from BacMet and RemeDB, reads were extracted, and kaiju was used. The downstream data analysis was performed in the R program and R studio (Version). Mainly microbiome and Phyloseq R package were used (McMurdie and Holmes 2013; Lahti and Shetty 2017). STAMP (v) (Parks et al., 2014) was used for differential abundance analysis, and plotting was done in GraphPad PRISM (8.0.2). The beta diversity analysis was performed using the Bray-Curtis distance method, and alpha diversity was analyzed using Shannon and chao1 indices. The data were normalized using Deseq2 normalization (Love et al., 2014). Further, ANOSIM and PERMANOVA using ADONIS-2 were used to interpret the statistics of beta diversity. For alpha diversity, we used the t-test and Wilcoxon test, depending upon the normality distribution of the data, which was performed in R using the Shapiro-Wilk normality test and Q-Q normalization plot. The differential abundance analysis in STAMP was performed for two groups using two-sided Welch’s t-test, White’s non-parametric test and ANOVA for multiple groups using Benjamini–Hochberg FDR correction test. The bacterial core was analysed using detection of 0.001 and prevalence of 50% and was plotted using Venny (v 2.1.0).

## 3. Results

### 3.1. Microbial community pattern

Using the Kaiju program, we used the nr_euk database to predict the microbial taxonomy pattern prevailing in the water hyacinth and corresponding environmental samples. Around 87% of the total reads were classified as Bacterial reads and ∼1 % as an Archaebacterial population. The details of the read distribution is stated in Supplementary Table 2. The phylum level analysis revealed *Proteobacteria* (Mean relative abundance (MRA) % in WH 2020 and 2022) (54.78±3.2% and 61.58±35%) as the predominant phylum, followed by *Bacteriodetes* (9.6±1.195% and 6.85±1.14%), *Actinobacteria* (6.87±1.13% and 7.1±1.19%), *Firmicutes* (4.04±0.56% and 3.8±1.05%), *Chloroflexi* (3.22±1.44% and 3.21±1.46%), *Verrucomicrobia* (3.71±0.5% and 1.98±0.5%), Cyanobacteria (2.61±1.8% and 0.86±0.26%), *Planctomycetes* (2.0±0.36% and 2.50±1.17%), *Acidobacteria* (1.75±0.42% and 2.16±1.31%), and *Spirochaetes* (1.36±0.4% and 0.46±0.1%) in WH (**Figure 1B**.). Among these dominant phyla, *Verrucomicrobia, Bacteriodetes*, and *Spirochaetes* were significantly higher in WH 2020 compared to 2022 (Welch’s t-test, Benjamini–Hochberg FDR, p<0.05, Supplementary Figure 1). Lake WH showed a significantly higher prevalence of *Actinobacteria* and S*pirochaetes* but a relatively lower *Proteobacterial* population than river WH (Welch’s t-test, Benjamini–Hochberg FDR, p<0.05, Supplementary Figure 2.). Furthermore, *Planctomycetes, Acidobacteria, Spirochaetes, Verrucomicrobia*, and *Chloroflexi* were differentially abundant in WH, whereas *Bacteroidetes* was higher in corresponding water samples in 2020 (White’s non-parametric test, Benjamini–Hochberg FDR, p<0.05, Supplementary Figure 3.). Similarly, *Firmicutes* were distinctly prevalent in WH of 2022 compared to the corresponding water samples (Welch’s t-test, Benjamini–Hochberg FDR, p<0.05, Supplementary Figure 4).

**Figure 1.**
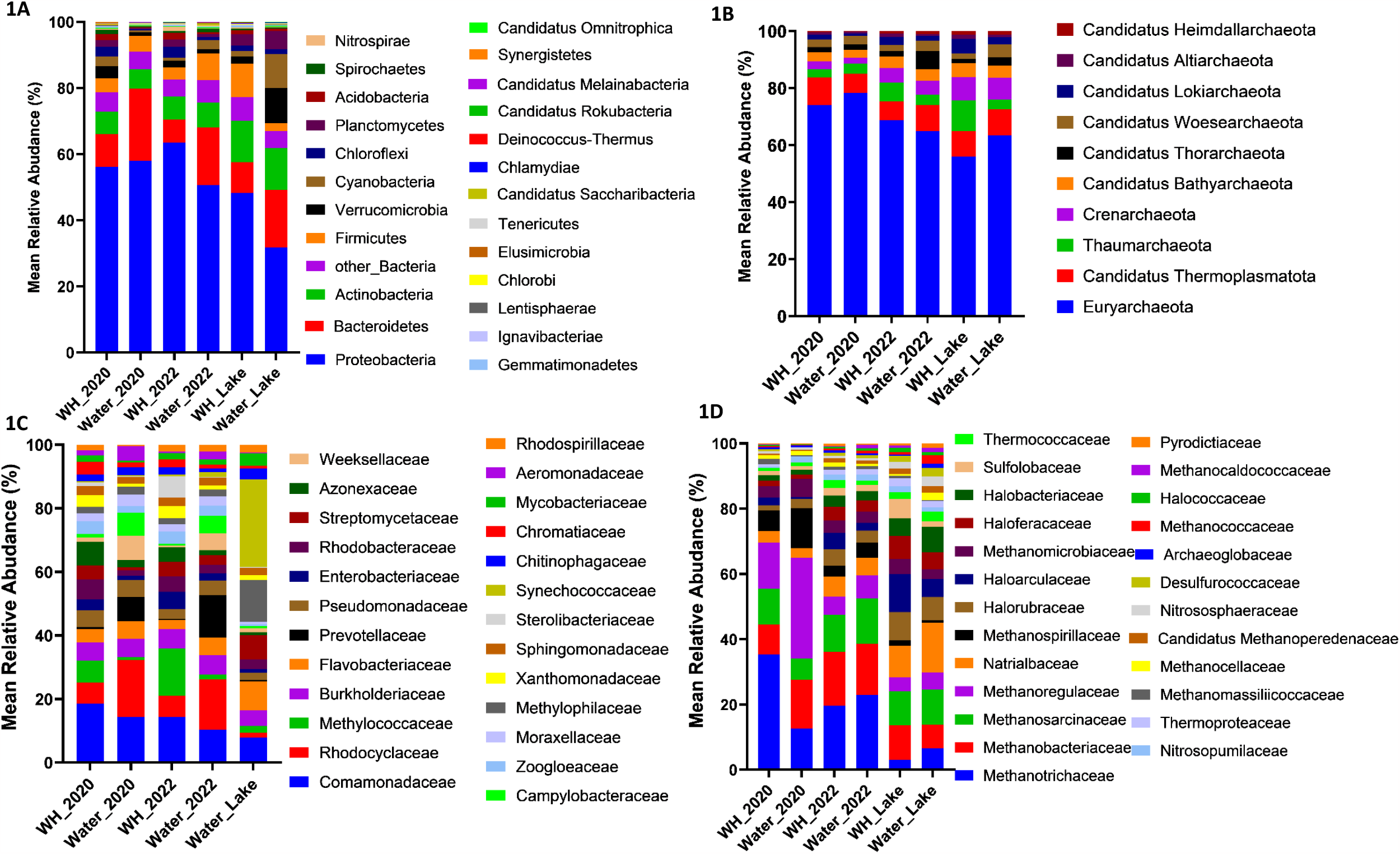
Microbial community pattern. The figure illustrates the percent mean relative abundance. of **1A** Bacterial phyla (top25), **1B**. Archaebacterial phylum, **1C**. Bacterial families (top 25), and **1D**. Archaebacterial families (top 25) in **WH_2020** (Water hyacinths from 2020 sampling), **WH_2022** (Water Hyacinths from 2022), **Water_2020** (Water samples from 2020), **Water_2022** (Water samples from 2022), **WH_Lake** (Water Hyaicnths from Pashan Lake), Water_Lake (Water from Pashan Lake).

*Betaproteobacteria, Gammaproteobacteria*, and *Alphaproteobacteria* comprise the most prevalent bacterial classes of the WH microbiome. *Alphaproteobacteria* were found to be significantly higher in 2022, whereas, *Verrucomicrobia, Bacteroidia, Spirochaetia*, and *Opitutae* were particularly predominant in WH 2020. The lake-river comparison of WH rhizobiome showed a significantly higher prevalence of *Cytophagia* and *Actinobacteria* in the lake and enrichment of *Betaproteobacteria* and *Deltaproteobacteria* in river WH. Furthermore, *Betaproteobacteria, Epsilonproteobacteria*, and *Flavobacteria* were found to be significantly enriched in water samples of 2020 (Supplementary Figure 5). Higher differential prevalence was observed at the lower level of the taxonomy. In WH vs Environment comparison, bacterial families (MRA %) *Comamonadaceae* (2-7%), *Rhodocyclaceae* (4 to 8%), *Campylobacteraceae* (1.3 to 3.7%), and *Burkholderiaceae* (2.4 to 2.8%) were amongst the most prevalent in water samples (Welch’s t-test, Benjamini–Hochberg FDR, p<0.05, Figure 1C.). Although not significant, *Prevotellaceae* members were more predominant in water samples than in WH (Average of 0.16% in WH_2020, 0.07% in WH_2022, 3.76% in Water_2020, and 5.19% in Water_2022) (Figure 1C.). We observed a higher and significant abundance of *Methylococcaceae, Rhodobacteriaceae, Azonexaceae*, and *Xanthomonadaceae* in WH_2022 (Welch’s t-test, Benjamini–Hochberg FDR, p<0.05). Further, Lake WH showed a significantly higher prevalence of *Streptomycetaceae, Nocardiaceae, Rhizobiaceae, Rhodobacteraceae, Mycobacteriaceae*, and *Phyllobacteriaceae*. In contrast, *Azonexaceae, Rhodocyclaceae, Comamonadaceae, Methylococcaceae, Chromatiaceae, Zoogleaceae, Desulfobacteriaceae, Desulfobulbaceae*, and *Desulfovibrionaceae* were more prominent in river WH (Welch’s t-test, Benjamini–Hochberg FDR, p<0.05).

The members of the phylum *Euryarchaeota* were the most prevalent (MRA 50% to 66%) in the WH of lake, river, and water samples. The other prominent phyla (MRA>1%) observed were *Candidatus Thermoplasmota, Thaumarchaeota, Crenarchaeota, Candidatus Bathyarchaeota, Candidatus Thorarchaeota, Candidatus Woesearchaeota, and Candidatus Lokiarchaeota. Methanomicrobia, Halobacteria* (Figure 1B.), *Methanomicrobia, Thermoplasmata, Thermoprotei*, and *Thermococci* were most abundant at the class level (Supplementary Figure 6). We observed a significantly differential prevalence of *Methanobacteria* (Welch’s t-test, Benjamini–Hochberg FDR, p<0.05) in 2020 water samples and a higher abundance of *Thermoplasmata* and *Halobacteria* (Welch’s t-test, Benjamini–Hochberg FDR, p<0.05) in WH20 and WH22, respectively. Further analysis revealed the predominance of *Methanotrichaceae* and *Natrialbaceae* in WH_2020 than in water samples. Although insignificant, we observed a differentially higher abundance of *Haloarculaceae, Sulfolobaceaeae, Methanocellaceae, Halorubraceae* (Welch’s t-test, Benjamini–Hochberg FDR, p<0.05), and *Halobacteriaceae* in WH_2022 than corresponding water samples Water_2022 (Figure 1D.).

## 3.2. Microbial community diversity of Water Hyacinth

Microbial community diversity of the water hyacinth rhizobiome was assessed using alpha diversity indices (chao1 and Shannon) and the Bray-Curtis dissimilarity distance method. Both analyses were performed to evaluate the influence of different sites or spatial effects (2020 and 2022), time (temporal 2020 vs 2022), environment (water), and aquatic bodies (river Vs Lake) on the WH rhizobiome. The microbial communities presented similar richness and diversity and no significant spatiotemporal differences (Figure 2, Supplementary Figure 7.). The diversity differences were observed in the WH rhizobiome from the lake and river. Bacterial and archaebacterial alpha diversity was higher in Lake WH compared to the WH of rivers (Figure 2A, Supplementary Figure 7). The comparison of the WH with their corresponding water samples reflected significantly higher (Wilcoxon test, p<0.05) microbial alpha diversity in WH, except in 2022 samples (Figure 2C.). The PCoA analysis using the Bray-Curtis distance method was performed to understand beta diversity. The WH microbiome varied significantly from their corresponding water samples with higher explanatory power and ANOSIM R-value (Figure 3C and 3D, Supplementary Figure 8A and B). Similarly, significantly higher microbial diversity differences were discerned after temporal and lake-river WH comparison (Figure 3B and 3A, Supplementary Figure 8C and D). Unlike the water hyacinth of 2020, significant site-based differences were observed in the river WH rhizobiome of 2022 (Supplementary Figure 9).

**Figure 2.**
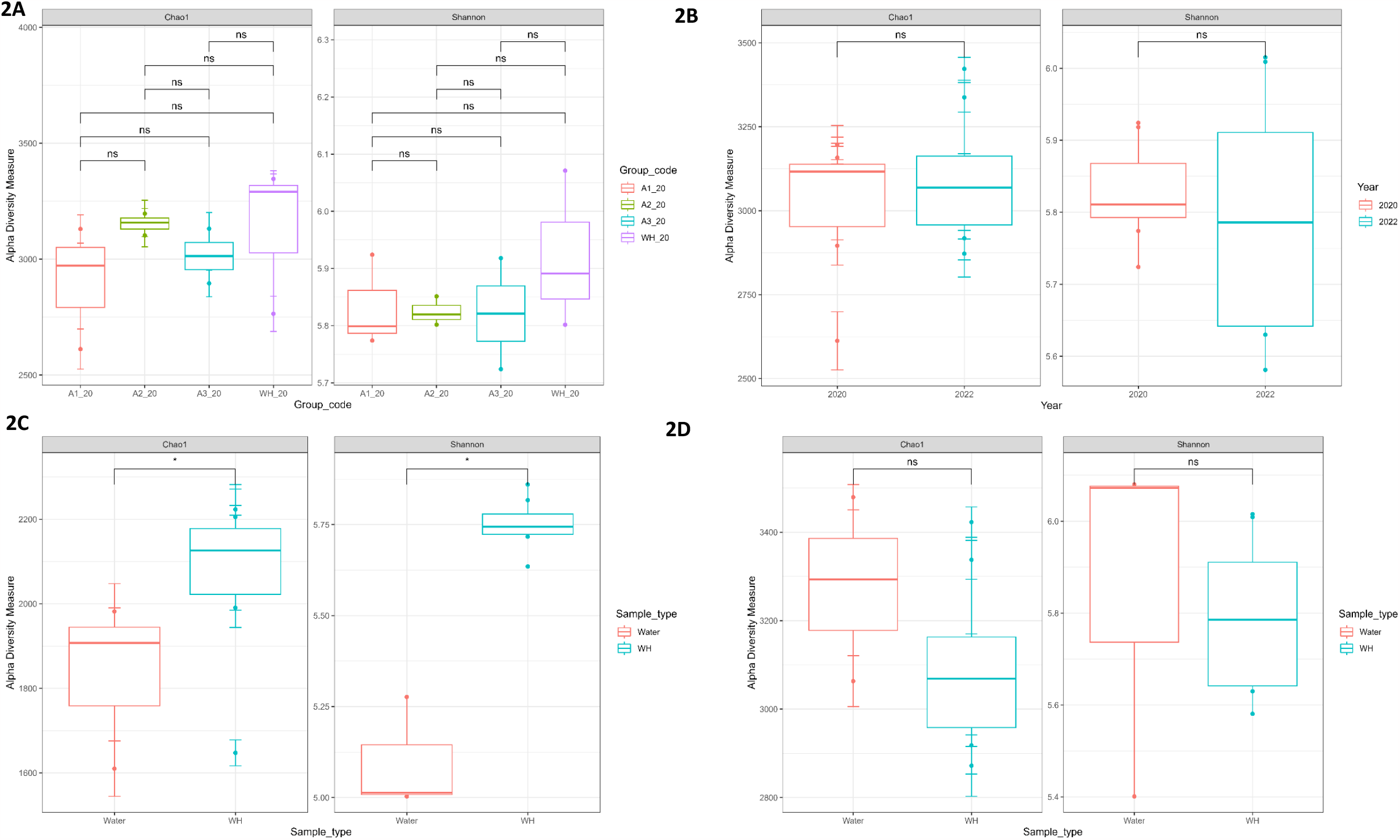
Bacterial Alpha Diversity. Box plots illustrates the alpha diversity indices chao1 and Shannon. of **2A** Alpha diversity between Lake WH and River WH (t-test, p>0.05), **2B**. Alpha diversity of River WH_2020 and WH_2022 (t-test, p>0.05), **2C**. Water hyaicnth of 2020 and Water samples 2020 (Wilcoxon test, p<0.05) and **2D**. Alpha diversity of WH_2022 and Water_2022 (t-test, p>0.05).

**Figure 3.**
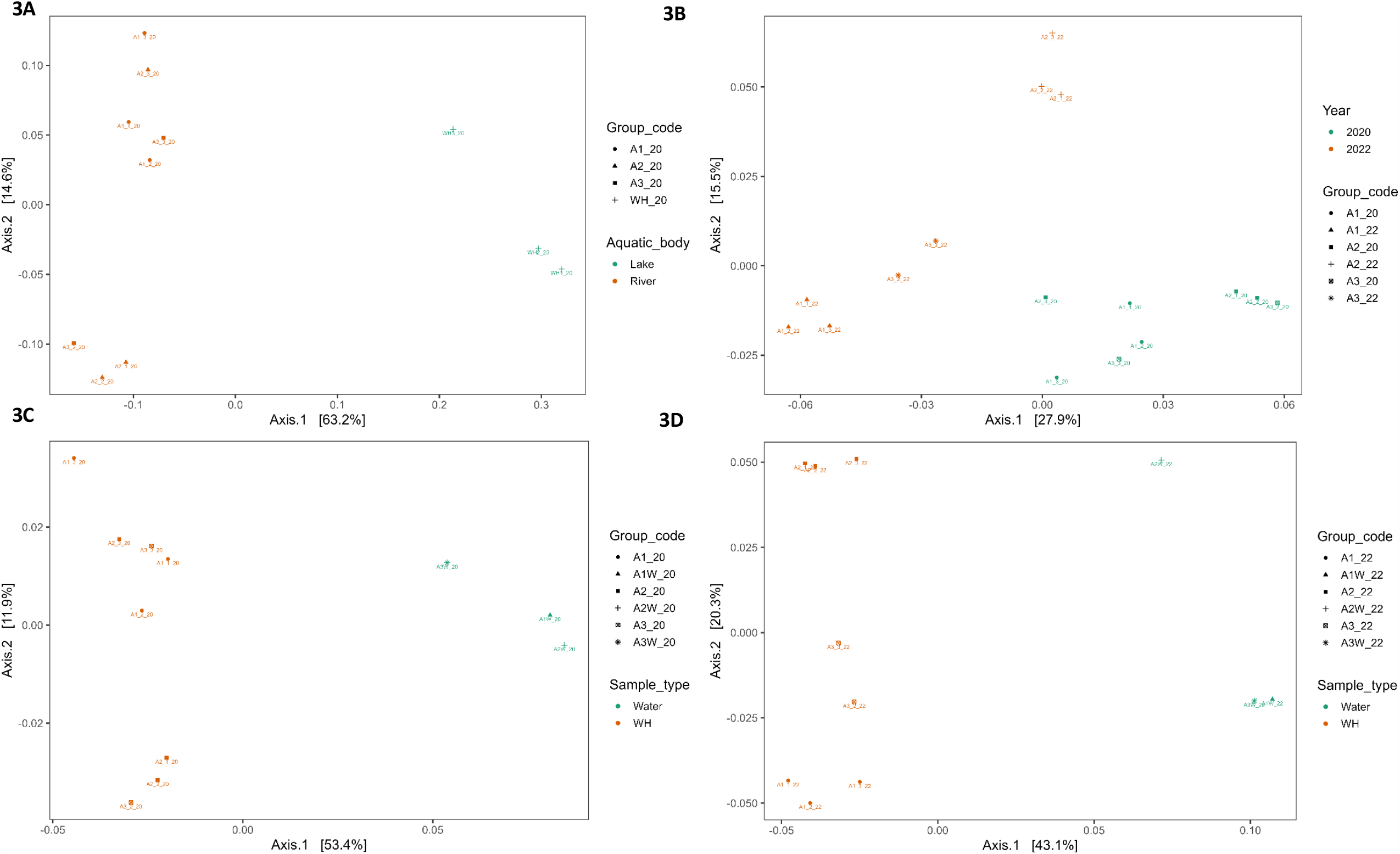
Bacterial Beta Diversity. PCoA plots using Bray-Curtis distance. **3A**. Between Lake WH and River WH (p<0.05, ANOSIM R=0.98, PERMANOVA ADONIS R2 =0.58, Beta-disper >0.05). **3B**. Between River WH_2020 vs WH_2022 (p<0.05, R=1, R2 =0.81, Beta-disper>0.05), **3C**. River Water_2020 and WH_2020 (p<0.05, R=1, R2 =0.52, Beta-disper>0.05), **3D**. River Water_2022 and WH_2022 (p<0.05, R=1, R2 =0.41, Beta-disper>0.05).

### 3.3. Core Microbiome

The core microbiome analysis was carried out at the family level of classification. For 2020 sampling, we observed that 45.5% of the core bacteria were found exclusively in WH than its corresponding water samples having ∼14.5% as unique and sharing 40% with the water hyacinth (Figure 4C.). Similar core uniqueness was observed for WH22, 42.6% absolute to WH, 19% to water, and the core sharing was 37.7% (Figure 4D). Comparing WH_2020 with WH_2022 revealed that most core bacteria were similar (47.7%), with the exclusive core as 24% and 27%, respectively (Figure 4B.). Although there was higher similarity in the core bacteria of Lake WH and River WH (37.8%), we also observed a highly unique (42.7%) core bacterial population in Lake (Figure 4A.).

**Figure 4.**
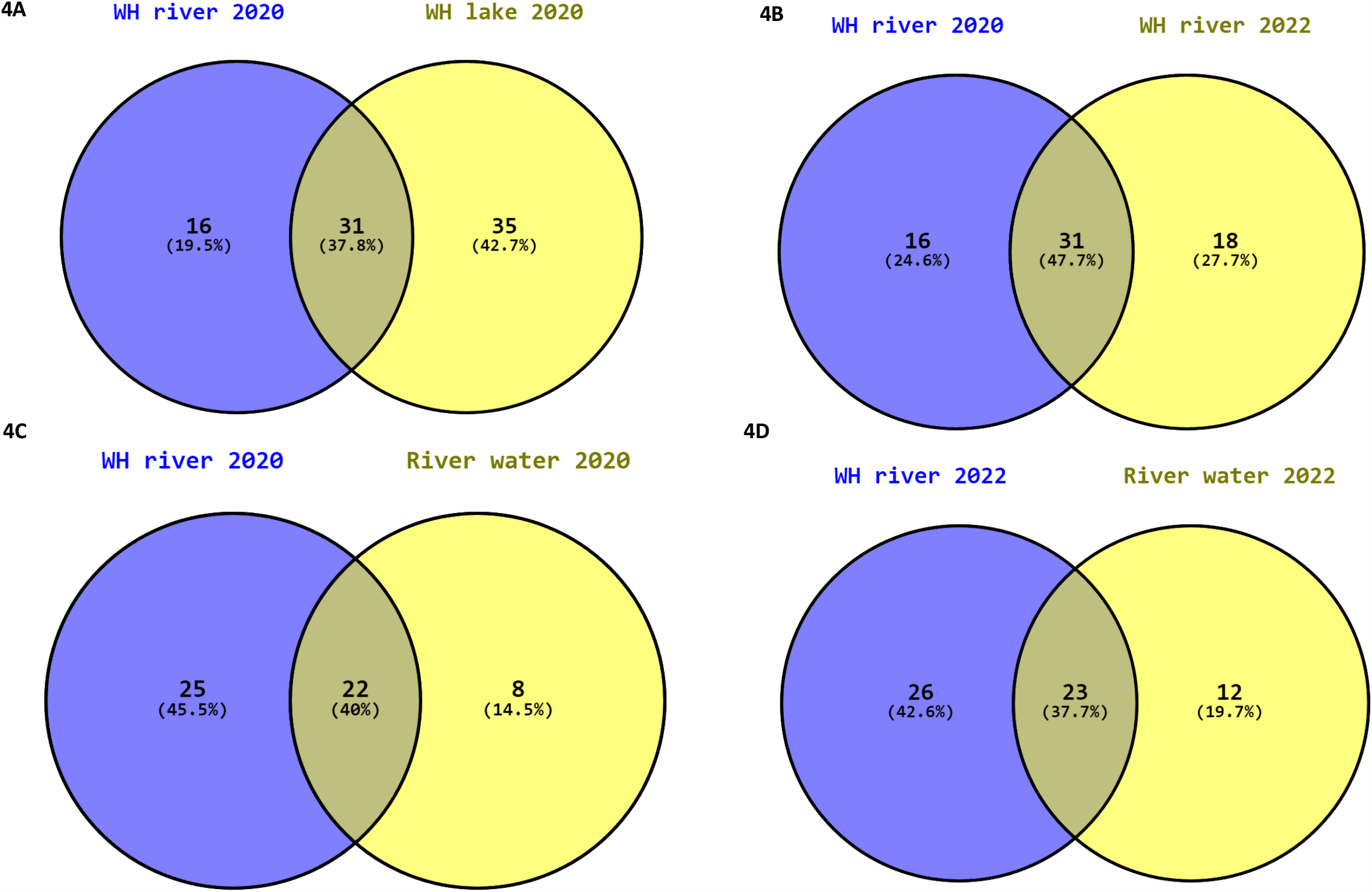
Bacterial core. Venn diagram for shared and exclusive bacterial families. The core was calculated using detection 0.001 and prevalence of 50%. **4A**. Core comparison between WH of lake and river, **4B**. Core comparison between WH of 2020 and 2022, **4C**. Core comparison of WH_2020 vs Water 2020, **4D**. Core comparison of WH_2022 and water 2022

### 3.4. Functional insights: Energy Metabolism and pollutant degrading enzymes

58% of the total bacterial and archaeal metagenomic reads were assigned to metabolic functions. Further, at the level 2 functional module of KEGG, 6% and 1.1% reads were assigned to Energy Metabolism and Xenobiotic metabolism, respectively. In energy metabolism, a total of 50% of the genes encoding for oxidative phosphorylation functions (52% in WH_2020, 52% in WH_2022, and 49.8% in Lake WH), followed by the metabolism of Nitrogen (21.2%), Methane (12.5%), Sulfur (7.48%), Carbon fixation (5.40%), and Photosynthesis (1.08%) (Figure 5A.). Among the xenobiotic metabolic functions, the majority of the reads were assigned for the metabolism of Nitrotoluene (28.4%), Benzoate (26.36%), Chlorocyclohexane and chlorobenzene (18.79%), Aminobenzoate (8.15%), Drug (4.02%), Atrazine (3.55%), Caprolactam (2.95%), Dioxin (2.13%), Xylene (1.69%), and Steroid (1.6%) (Figure 5B). We further predicted the Pollutant Degrading Enzymes (PDE for hydrocarbon, plastic, and dye degradation) in the WH microbiome using RemeDB. The Catalase peroxidase (10%) was the most abundant PDE in water hyacinth rhizobiome, followed by Protocatechuate 4,5-dioxygenase (5.5%), Chemotaxis response regulator protein-glutamate methylesterase (5.38%), 2-nitropropane dioxygenase (5.17%%), and Glyoxalase/bleomycin resistance (3.16%) (Figure 5C). The taxonomic classification of the PDEs revealed the predominance of *Proteobacterial* classes, with *Betaproteobacteria* (64.4%), *Gammaproteobacteria* (11.48%), and *Alphaproteobacteria* (7.12%) being more prevalent. The other prominent bacterial types (>0.2%) contributing to PDEs in WH rhizobiome were *Actinobacteria, Deltaproteobacteria, Falvobacteria, Cytophagia, Hydrogenophilalia*, and *Bacteroidia*. Further, at the genus level, we observed a higher abundance (>2%) of *Hydrogenophaga, Ideonella, Rubrivivax*, and *Dechloromonas*. The other numerically prominent members (>1%) were *Acidovorax, Methylomonas, Pseudomonas, Aquabcterium, Thauera*, and *Sphaerotilus* (Supplementary Figure 10 and 11).

**Figure 5.**
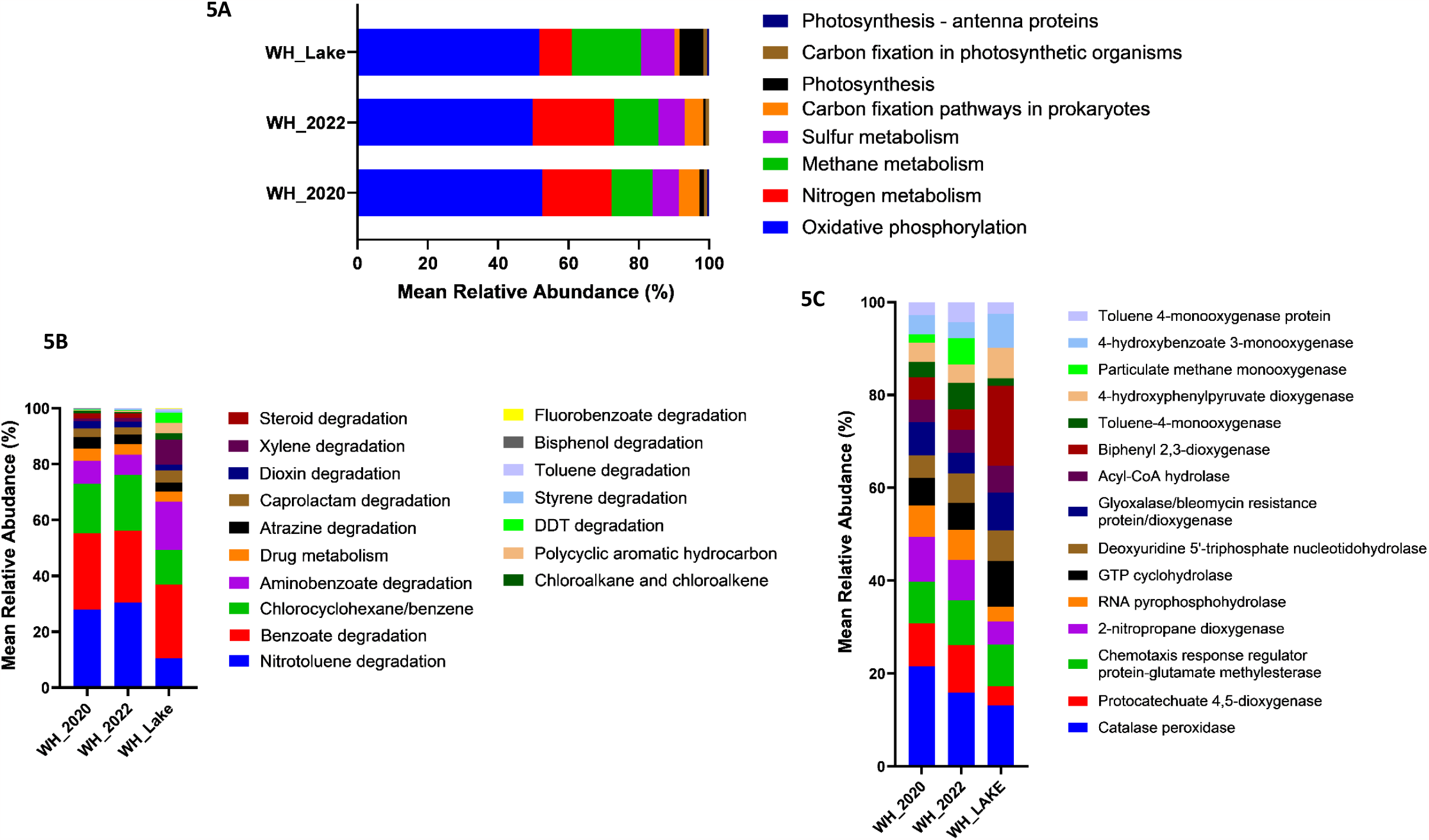
Energy and Bioremediation metabolism of WH microbiome. The stacked bar plot illustrates **5A**. Prevalence of Energy metabolism pathways, **5B**. Xenobiotic metabolism, **5C**. Top 15 Pollutant degrading enzymes in WH microbiome using RemeDB. The gene level information is provided in supplementary tables

### 3.5. Biocides and Metal resistance genes (MRGs)

The BacMet database analysis revealed 341 genes encoding antibacterial biocide and metal resistance in the WH microbiome. Altogether, we observed higher resistance to Biocides and other compounds (Percent abundance, WH_2020 (23.97%), WH_2022 (24.27), WH_LAKE (21.13%)), followed by Metals and other compounds (13.21%, 15.6%, 21.77%), Copper (14%, 9%, 17%), Multi-metal resistance (11%, 12%, 13%), Arsenic (11%, 13%, 12%), Mercury (8%, 7%, 3%), Chromium (4%, 5%, 4%), and other heavy metals (Supplementary Figure **12**). At the gene level, ruv B, a DNA helicase-mediated resistance gene, were highly prevalent (9%) in WH rhizobiome, followed by *sodB* (8%), *dpsA* (7%), *merA, actP, cop, acr3, chrA, mdeA, arsC, copR, abeS, pcm, copF, abeS, pcm*, and *copF* (>1%) (Figure 6A.). Further, the majority of the genes were classified as *Methylomonas* (1.99%), *Dechloromonas* (1.8%), *Hydrogenophaga* (1.56%), *Nitrospira* (1.02%), *Ideonella* (0.87%), *Pseudomonas* (0.83%), *Rubrivivax* (0.82%), *Planktothricoides* (0.72%), and *Acidovorax* (0.6%) (Figure 6B.).

**Figure 6.**
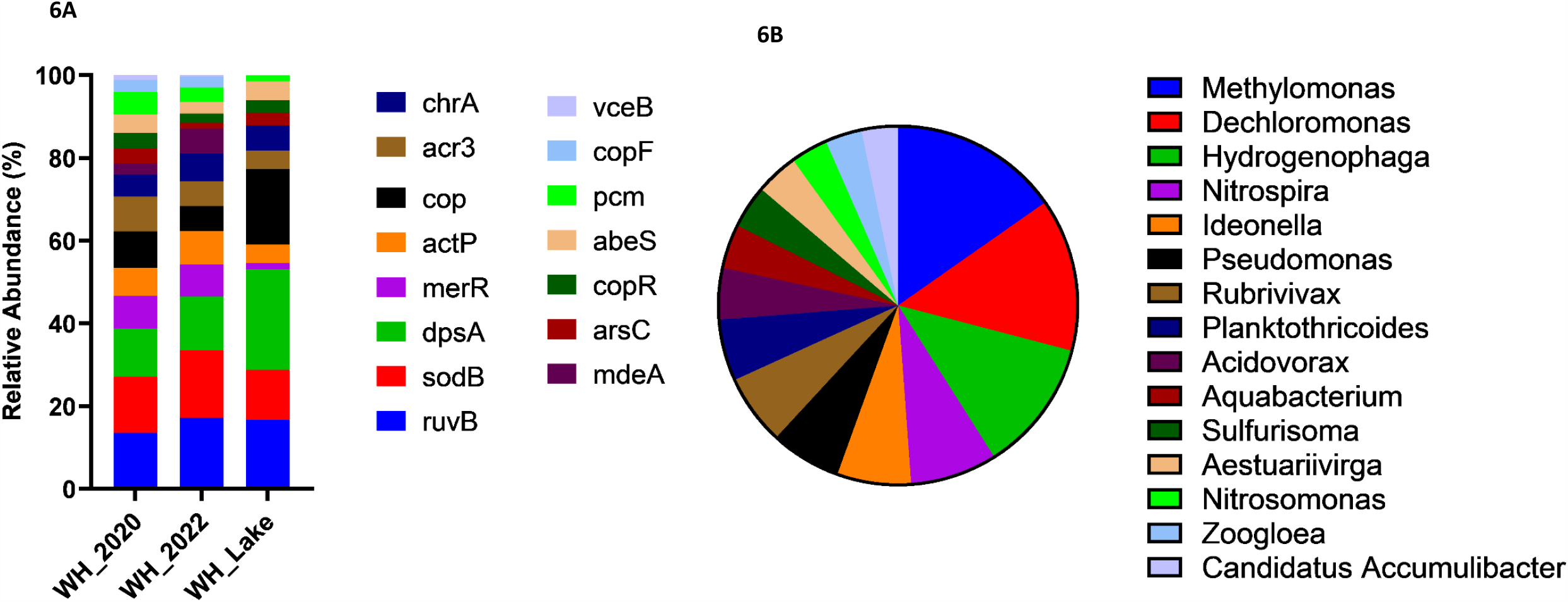
Biocides, Metal Resistance, and bacterial classification. 6A. Top 15 Biocide and Metal resistance genes in WH microbiome, 6B. Prevalent bacterial genus encoding Metal and Biocide resistance.

## 4. Discussion

Here, we investigated the diversity (spatial and temporal) and functional potentials of the water hyacinth root-associated microbiome in rivers and a lake in Pune city, India. We observed substantial differences in the WH microbial diversity by the type of aquatic bodies (River and Lake), with its surrounding environment (water) and on a temporal scale. Moreover, our analysis of WH rhizobiome also indicated their remediating ability and tolerance to biocides and metals. We observed diverse bacterial communities associated with the river water hyacinth, particularly *Proteobacteria* being the most dominant, followed by *Bacteriodetes, Actinobacteria, Firmicutes, Chloroflexi*, and *Verrucomicrobia* (Figure 1.). A previous study on *E. crassipes* from Hindon River, India, by Sharma et al. 2021, observed similar numerically abundant bacterial taxa. In comparison to the WH from wetlands (Ávila et al., 2019), we observed a relatively higher proportion of *Bacteroidetes* members (10%) in the river WH. Additionally, a higher prevalence of *Euryarchaeota* was observed in river WH, whereas, in wetlands, the *Woesearchaeota* phylum was more dominant (Figure 1.). Overall, these differing observations can be attributed to the higher occurrence of these taxa in the rivers of Pune city (Yadav et al., 2021), the effect of geography, and the difference in the type of aquatic ecosystems (Ávila et al., 2019).

### 4.1. Aquatic bodies (Habitats) drives the variance in the rhizobiome of Water Hyacinth

Contrasting observations were noted in the microbiome of WH from Lake, where a significantly higher enrichment of *Actinobacteria* and *Spirochaetes* and a lower abundance of *Proteobacteria* was observed than in the river WH (Welch’s t-test, Benjamini–Hochberg FDR, p<0.05). Similar observations were recorded at the lower level of taxonomy, where bacterial families such as *Streptomycetaceae, Nocardiaceae, Rhizobiaceae, Rhodobacteraceae, Mycobacteriaceae*, and *Phyllobacteriaceae* were significantly (Welch’s t-test, Benjamini–Hochberg FDR, p<0.05) enriched in Lake WH compared to river WH showing the differential abundance of *Azonexaceae, Rhodocyclaceae, Comamonadaceae, Methylococcaceae, Chromatiaceae, Zoogleaceae, Desulfobacteriaceae, Desulfobulbaceae*, and *Desulfovibrionaceae*. Although insignificant, we observed more archaebacterial lineages such as *Natrialbaceae, Methanosarcinaceae*, and *Haorubraceae* in the Lake WH than in the river WH (Figure 1D.). Moreover, the Lake WH microbiome exhibited higher microbial alpha diversity than the river WH. Additionally, we observed significantly distinct clustering of microbiome of Lake WH and river WH upon beta diversity estimation. The higher ANOSIM R-value and PERMANOVA R^2^ value using the ADONIS-2 test (R-value and R^2^ value for bacteria and Archaea (0.98, 0.57%) and (0.94, 0.36%), respectively) suggested aquatic bodies as a decisive explanatory variable for microbial shaping of water hyacinth (Figure 3A. and Supplementary Figure 8C.). Rivers and lakes are distinct habitats with different ecological processes, anthropogenic impacts, and multiple biotic interactions that determine the microbiological community (Tang et al., 2020). The core microbiome analysis further substantiated the influence of habitats. Although river and Lake WH shared 37 % of their core bacterial population, around 43% of the core bacteria were exclusive to Lake WH (Figure 4A.). Among these core bacteria, some of the dominant bacterial families unique to Lake WH were *Zhaonellaceae, Weeksellaceae, Synechococcaceae, Rhodospirillaceae, Pseudonocardiaceae, Cytophagaceae*, and others. Overall, these observations suggested the prominent influence of the aquatic bodies in determining the water hyacinth’s microbial community.

### 4.2. Temporal variations visible in transient microbiome

Upon comparison of the River WH microbiome in the year 2020 (WH_2020) and 2022 (WH2022), we observed differential enrichment of microbial taxa. The bacterial phylum *Verrucomicrobia, Bacteriodetes*, and *Spirochaetes* were significantly higher in WH 2020 compared to 2022. Similarly, the bacterial families such as *Prevotellaceae, Bacteroidaceae, Spirochaetaceae, Desulfovibrionaceae*, and *Desulfobacteraceae* were significantly higher in WH_2020 and *Gallionellaceae* and *Hyphomicrobiaceae* in WH_2022. We did not observe differential preference in the archaebacterial population; however, we observed a higher prevalence of *Methanotrichaceae* in WH_2020 samples (Mean abundance of 14% vs 8% in WH_2022). Unlike the lake WH microbiome, the core microbiome in the river on a temporal scale was much conserved (48% of the core was common), with only 24% to 27% being unique, which can be attributed to the environmental habitat (Figure 4B.). We did not observe a significant temporal influence on the overall alpha diversity but observed significant beta diversity differences in the microbiome of WH_2020 and 2022. The ANOSIM and PERMANOVA revealed significantly higher differences with R-values of 1.0 for bacteria and 0.41 for archaebacteria suggesting distinct microbial cluster formation and R^2^ values of 0.81% for bacteria and 0.17% for archaea, indicating that most of the observed variations are as a result of temporal factor (Figure 3B, Supplementary Figure 8D.). These results indicated changes in the overall microbiome of water hyacinth in rivers across time. Despite the observed variations, most of the core microbiomes in the river water hyacinth of 2020 and 2022 were conserved (Figure 4B.). Thus these variations could be associated with the transient flora and surrounding water.

### 4.3. WH microbiome is distinct than its surrounding water

The differential enrichment of microbial population in water hyacinth was quite visible compared to their corresponding water samples. *Bacteroidetes* and *Firmicutes* were significantly higher in the water samples of the year 2020 and 2022, respectively, whereas the WH harboured significantly higher members of *Planctomycetes, Acidobacteria, Spirochaetes, Verrucomicrobia*, and *Chloroflexi* (Supplementary Figure 4 and 5.). *Bacteroidetes* and *Firmicutes* are highly predominant in rivers from urban cities (Lin et al., 2019; Yadav et al., 2021). Along with freshwater bacterial groups such as *Flavobacteraceae*, we observed a significantly higher abundance of gut-related and pathogen-containing groups such as *Campylobacteraceae, Burkholderiaceae*, and *Prevotellaceae* in water samples than in water hyacinth (Bernardet and Nakagawa et al., 2006; Magana-Arachchi & Wanigatunge, 2020) (Supplementary Figure 12). Further, the organisms reported for bioremediation (Loredanan et al., 2017; Zhang et al., 2021; Yadav et al., 2020), such as *Rhodobacteriaceae, Azonexaceae, Xanthomonadaceae, Rhodobacteraceae, Azonexaceae*, and few others were numerically more abundant in the WH than water (Supplementary Figure 12). Also, significantly higher occurrences of methanogens and methanotrophs such as *Methylococcaceae, Methanotrichaceae*, and *Methanocellaceae* were observed in WH than in water (Supplementary Figure 12). Interestingly, most of the core microbiome was shared between water hyacinth and water; however, WH still had unique 45% and 42% unique core bacterial families in the years 2020 and 2022, respectively (Figure 4C and 4D). These observations indicate that WH has a stable core microbiome which is probably maintained over time irrespective of its environment and is also supported by the earlier observations with temporal and aquatic body comparison. Furthermore, from the diversity perspective, we observed significantly higher alpha diversity in the WH microbiome than in water samples. Upon beta diversity estimation using the Bray-Curtis distance measure, we observed robust clustering and variance explanation value (ANOSIM R of 0.46 to 1.0 and R^2^ of 0.18 to 0.52), indicating significantly higher diversity differences between the water and WH microbiome (Figure 3C and 3D.). Overall, these observations indicated that probably water hyacinth maintains probably selectivity or vertical transfer of microbiome and harbour significantly different diversity from their surrounding water environment. In future, it will be interesting to understand detailed molecular insights, including the chemical signalling and interactions existing between the roots and surrounding microbes in water to maintain such selectivity.

### 4.4. Energy metabolism and microbial-associated bioremediation

Previously, Ávila et al., 2021, reported root-associated methanogenesis and oxidation as primary energy-deriving metabolic functions in water hyacinth. Our study observed oxidative phosphorylation (aerobic process) and nitrogen metabolism as the dominant energy metabolism functions, followed by methane, sulphur, and carbon metabolism. The urban aquatic bodies are usually influx by a large amount of nitrogen inputs amid higher anthropogenic activities (Yu et al., 2020). The presence of nitrification genes (amoA, B, C), denitrification genes (nir, nar, nor, nos, nap), and core-forming bacterial groups such as *Acetobacteraceae, Bradyrhizobiaceae*, and *Rhizobiaceae* substantiated the higher prevalence of nitrogen metabolism in WH of urban aquatic bodies. The methane metabolism was quite prevalent in Lake WH (19%) as compared to river WH (11%), whereas nitrogen metabolism was more prevalent in river WH (23%) than in Lake WH (9%). Studies (Xiao et al., 2017; Hilt et al., 2022, Kosten et al., 2016) have reported that methane concentrations and fluxes in aquatic bodies are highest in the vegetated zones by macrophytes. The core microbiome of the WH consisted of the methane-metabolizing archaebacterial groups (Liu, 2010; Costa et al., 2019) such as *Methanobacteriaceae, Methanobacteriaceae, Methanocellaceae, Methanomicrobiaceae, Methanoregulaceae, Methanosarcinaceae, Methanospirillaceae, Methanotrichaceae* suggesting that the root-associated archaebacteria mainly drive methane metabolism in WH. Further, the abundance of bacterial groups (core microbiome) such as *Desulfobacterales, Desulfovibrionales*, and *Desulfuromonadales* can be linked to sulfur metabolism in water hyacinths. Such sulphate-reducing bacteria are reported to be associated with water hyacinth (Achá et al., 2005). Several studies have reported the use of WH to reduce water pollutants such as organic dyes, metals, and insecticides, mainly attributing their removal to the functional groups (—PO4, C=O, and C—H) on the root surfaces (Madikizela et al., 2021; Delgado et al., 1993). A study by Shehzadi et al., 2016, also reported the enhancement in the remediation of pollutants with the associated endophytic bacteria of *E*.*crassipes*, facilitating bioremediation. Certain microbes extract energy from metabolizing xenobiotic compounds, especially in this study’s polluted river ecosystem. 1.1% of the total reads were assigned to xenobiotic degradation and metabolism, whereas top reads were assigned to Nitrotoluene and Benzoate degradation. Our previous study (Yadav et al., 2020) from the same river sites revealed similar enrichment of genes in sediments involved in the degradation of these compounds. The WH in the present study are from polluted rivers and are often exposed to pollutants, which might play a role in determining the microbial structure of WH. Further, to understand other remediating functionalities, we used the RemeDB database to mine for pollutant-degrading enzymes (PDEs) in the microbial population associated with water hyacinth. We observed over 140 PDEs in the WH microbiome with dye, hydrocarbon, and plastic degrading abilities. Various dye-degrading enzymes such as Catalase peroxidase and other peroxidases, Catechol 1,2-dioxygenase; hydrocarbon utilizing enzymes such as monooxygenases and dioxygenases; and plastic degrading enzymes such as catalase, phenylacetate dehydrogenase, and various hydrolases were detected (Yadav et al., 2020; Mooney et al., 2006; 21. Sankara Subramanian et al., 2019). The taxonomic classification of the PDEs predicted a higher prevalence of *Hydrogenophaga, Ideonella, Rubrivivax, Dechloromonas, Thauera, Pseudomonas* and several others that are already reported for degradation of xenobiotic compounds such as aromatic hydrocarbons and plastic degradation (Supplementary Figure 11) (Palm et al., 2019; Dubbels et al., 2009; and Fan et al., 2019).

### 4.5. Metal removal by water hyacinth or WH’s root-associated microbiome?

Several studies have reported hyacinth for efficiently removing metals and biocides (Delgado et al., 1993; Polprasert & Khatiwada, 1998; Rezania et al., 2015; Nor, 1994). Recent studies have also reported enhancing metal and biocides removal by plant-associated bacteria (Anudechakul et al., 2015; Mahfooz et al., 2021; Kularatne et al., 2009). The plant roots are hot spots for biodiversity, providing a favourable ecological niche for microbe-plant and microbe-microbe interactions (Sharma et al., 2021). The roots of water hyacinth absorb heavy metals and biocides, which can be essential to the associated microbes at a lower concentration. However, at high concentration these could harm microbes. Microorganisms have thus developed various mechanisms to tolerate these metals and biocides via different mechanisms, such as inactivation or transformation of metals, impermeability to metals, and alteration of sites (Trevors et al., 1985). The analysis revealed that higher resistance to biocides and metals is associated with the WH microbiome. Higher resistance to metals such as Copper, Arsenic, Mercury, Chromium, Antimony, Lead, and Zinc was observed. Furthermore, we observed higher *ruvB* and *sodB*-mediated resistance at the gene level. The RuvB is an ATP-dependent helicase which can repair the DNA damage caused by toxic metals (BacMET, Miranda et al., 2005). The *sodB* encodes superoxide dismutases that are reported to protect bacterial cells against metal oxides (BacMET, Bébien et al., 2002). In addition to these two genes, we observed around 340 metal and biocide resistance genes in WH rhizobiome, such as *dps A* for hydrogen peroxide, *mer R* for mercury, *cop* for copper resistance, act gene for multiple metals and biocides, *acr* for Arsenic and several others (BacMET). The taxonomic classification revealed bacterial genera such as *Methylomonas, Dechloromonas, Hydrogenophaga, Nitrospira, Pseudomonas, Ideonella*, and others that contributed to metal and biocide resistance. Most of these, such as *Methylomonas, Hydrogenophaga, Ideonella, Pseudomonas*, and *Thauera*, constitute the core microbiome of WH, thus indicating that metal removal by WH is greatly facilitated by its microbiome. These WH-associated metal and biocide-resisting microbiomes could also have biotechnological applications for controlling toxic metals in various environments. On the other hand, these traits could also lead to the spread and dissemination of antibiotic resistance. The evolution of antibiotic resistance is not only governed by antibiotics but also by other factors, such as metals, as they possess a common mode of action and thus could be co-selected (Li et al., 2017).

## 5. Conclusion

To conclude, we observed diverse microbial communities associated with water hyacinth roots. The microbial diversity of the water hyacinth was significantly affected by the habitats (type of aquatic bodies), revealed by distinct and non-overlapping microbial cluster formation. Further, temporal variations were also observed in the water hyacinth microbiome that could be associated with the transient microbial taxa, as most of the core taxa were conserved. These observations are also supported by the fact that most of the core taxa of the WH were unique compared to its surrounding water and significant differences between the water and WH microbiome. We observed significantly differentially prevalent taxa in WH compared to water, such as methane and sulfur metabolizing organisms. Collectively, these observations suggested the selectivity of WH in maintaining its microbiome. Further, in line with the previous studies, we observed methane metabolism as one of the significant energy-generating mechanisms in water hyacinth. In addition to methane metabolism, we also revealed a higher prevalence of sulfur-metabolizing microbial taxa constituting the core microbiome of WH. Furthermore, the analysis revealed various xenobiotic remediation genes and pollutant-degrading enzymes in the WH microbiome. The detection of a higher abundance of metal resistance and biocide genes in the WH microbiome suggested the role of microbes in the remediation of pollutants by water hyacinth. The present study provided detailed information about the overall microbiome of water hyacinth, the factors affecting diversity, and the functionality of the WH microbiome. The present study provides the baseline information for conducting future research that shall be directed towards understanding the selectivity of the WH in maintaining its microbiome and co-occurrence of metal and antibiotic resistance in water hyacinth.

## 6. Acknowledgements

Authors are grateful to the Director, CSIR-NCL for providing facilities, infrastructure and support. RKY would also acknowledge University Grants Commission (UGC) for fellowship. We are thankful to Mr Hardik Chavda for his assistance in sampling in the year 2020.

## 7. Authors contribution

Rakeshkumar Yadav-Data curation, Formal analysis, Investigation, Methodology, Writing; Vinay Rajput-Formal analysis; Mahesh Dharne-, Conceptualization, Supervision, Resources, Writing - review & editing.

