## Supplementary information for "Water Hyacinth microbiome: metagenomic cues from environment and functionality in urban aquatic bodies"

**Supplementary information for research article entitled “Water Hyacinth microbiome: metagenomic cues from environment and functionality in urban aquatic bodies”.**

**Supplementary Table 1.** The following table provides details of water hyacinth and water samples. WH= Water Hyacinth. Here, Mula and Ramnadi are rivers and Pashan is a lake.

| **Sr. No.** | **Sample codes** | **Group code** | **Sample type** | **Aquatic body** | **Broad codes** | **River/Lake** | **Year of sampling** |
|---|---|---|---|---|---|---|---|
| 1 | A1_1_20 | A1_20 | WH | River | WH_2020 | Mula | 2020 |
| 2 | A1_2_20 | A1_20 | WH | River | WH_2020 | Mula | 2020 |
| 3 | A1_3_20 | A1_20 | WH | River | WH_2020 | Mula | 2020 |
| 4 | A2_1_20 | A2_20 | WH | River | WH_2020 | Ramandi | 2020 |
| 5 | A2_2_20 | A2_20 | WH | River | WH_2020 | Ramnadi | 2020 |
| 6 | A2_3_20 | A2_20 | WH | River | WH_2020 | Ramnadi | 2020 |
| 7 | A3_2_20 | A3_20 | WH | River | WH_2020 | Mula | 2020 |
| 8 | A3_3_20 | A3_20 | WH | River | WH_2020 | Mula | 2020 |
| 9 | A1W_20 | A1W_20 | Water | River | Water_2020 | Mula | 2020 |
| 10 | A2W_20 | A2W_20 | Water | River | Water_2020 | Ramnadi | 2020 |
| 11 | A3W_20 | A3W_20 | Water | River | Water_2020 | Mula | 2020 |
| 12 | A1_1_22 | A1_22 | WH | River | WH_2022 | Mula | 2022 |
| 13 | A1_2_22 | A1_22 | WH | River | WH_2022 | Mula | 2022 |
| 14 | A1_3_22 | A1_22 | WH | River | WH_2022 | Mula | 2022 |
| 15 | A2_1_22 | A2_22 | WH | River | WH_2022 | Ramandi | 2022 |
| 16 | A2_2_22 | A2_22 | WH | River | WH_2022 | Ramnadi | 2022 |
| 17 | A2_3_22 | A2_22 | WH | River | WH_2022 | Ramnadi | 2022 |
| 18 | A3_2_22 | A3_22 | WH | River | WH_2022 | Mula | 2022 |
| 19 | A3_3_22 | A3_22 | WH | River | WH_2022 | Mula | 2022 |
| 20 | A1W_22 | A1W_22 | Water | River | Water_2022 | Mula | 2022 |
| 21 | A2W_22 | A2W_22 | Water | River | Water_2022 | Ramnadi | 2022 |
| 22 | A3W_22 | A3W_22 | Water | River | Water_2022 | Mula | 2022 |
| 23 | WH1_20 | WH_20 | WH | Lake | WH_Lake | Pashan | 2020 |
| 24 | WH2_20 | WH_20 | WH | Lake | WH_Lake | Pashan | 2020 |
| 25 | WH3_20 | WH_20 | WH | Lake | WH_Lake | Pashan | 2020 |
| 26 | WHW_20 | WHW_20 | Water | Lake | Water_Lake | Pashan | 2020 |

**Supplementary Table 2.** The following table provide the details of total classified and unclassified reads after taxonomic classification by Kaiju using nr_euk database.

| Sr. No. | Sample codes | Classified | Unclassified |
|---|---|---|---|
| 1 | A1W_20 | 114333 | 13844 |
| 2 | A1W_22 | 49378 | 71167 |
| 3 | A1_1_20 | 43854 | 27369 |
| 4 | A1_1_22 | 73377 | 57338 |
| 5 | A1_2_20 | 51899 | 35879 |
| 6 | A1_2_22 | 42339 | 33142 |
| 7 | A1_3_20 | 17695 | 8585 |
| 8 | A1_3_22 | 48764 | 76276 |
| 9 | A2W_20 | 45339 | 4443 |
| 10 | A2W_22 | 232396 | 123263 |

| | | | |
|---|---|---|---|
| 11 | A2_1_20 | 63395 | 52819 |
| 12 | A2_1_22 | 147821 | 64386 |
| 13 | A2_2_20 | 54521 | 37168 |
| 14 | A2_2_22 | 64763 | 40147 |
| 15 | A2_3_20 | 73577 | 52396 |
| 16 | A2_3_22 | 41730 | 83081 |
| 17 | A3W_20 | 68296 | 11797 |
| 18 | A3W_22 | 83954 | 113734 |
| 19 | A3_2_20 | 43969 | 15951 |
| 20 | A3_2_22 | 164639 | 250970 |
| 21 | A3_3_20 | 38670 | 26902 |
| 22 | A3_3_22 | 68929 | 85046 |
| 23 | WH1_20 | 23889 | 195162 |
| 24 | WH2_20 | 44763 | 352037 |
| 25 | WH3_20 | 99416 | 395122 |
| 26 | WHW_20 | 145425 | 122904 |

**Supplementary Figure 2.** Extended error bar plot showing differentially abundant bacterial phylum (Welch's t-test, Benjamini–Hochberg FDR, $p<0.05$) between WH_2020 and WH_2022.

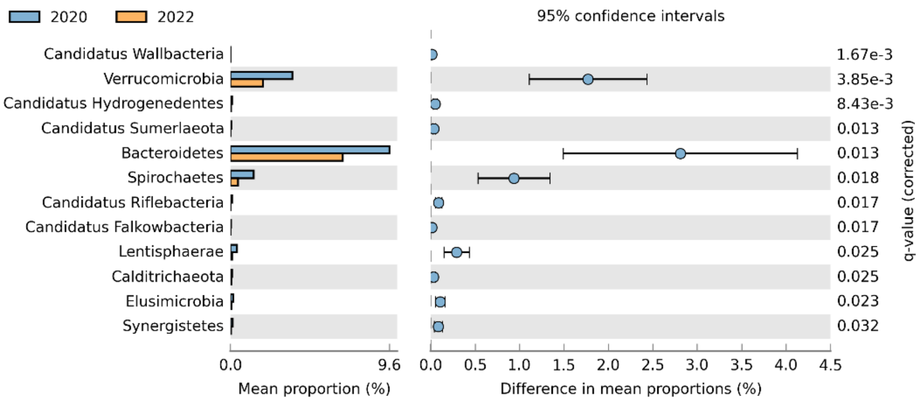


**Supplementary Figure 2.** Extended error bar plot showing differentially abundant bacterial phylum (Welch's t-test, Benjamini–Hochberg FDR, $p<0.05$) between Lake and River.

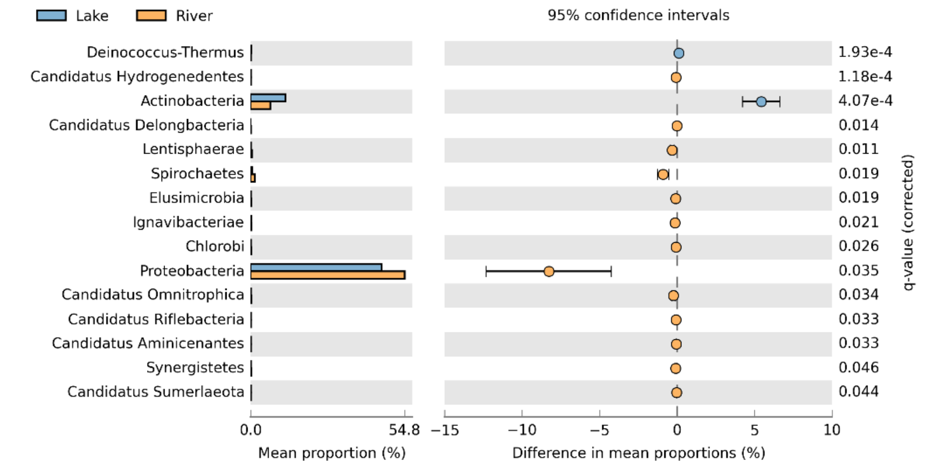


**Supplementary Figure 3.** Extended error bar plot showing differentially abundant bacterial phylum (White's non-parametric test, Benjamini–Hochberg FDR, p<0.05) between Water_2020 and WH_2020.

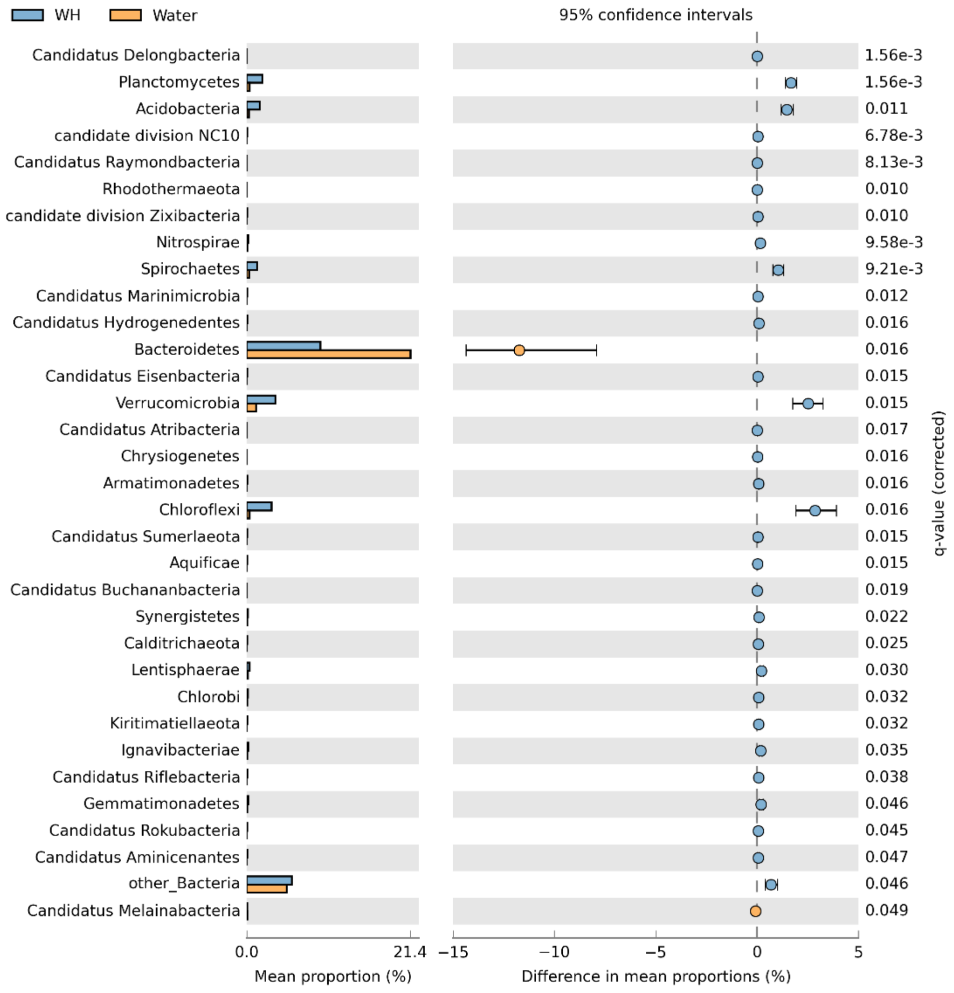


**Supplementary Figure 4.** Extended error bar plot showing differentially abundant bacterial phylum (Welch's t-test, Benjamini–Hochberg FDR, p<0.05) between Water_2022 and WH_2022.

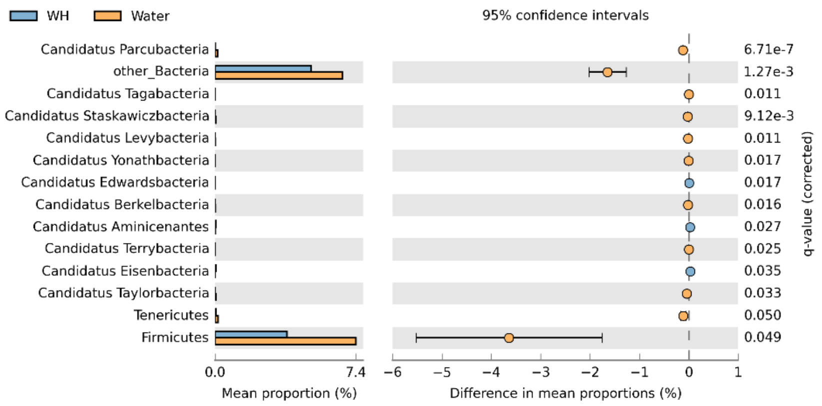


**Supplementary Figure 5.** Extended error bar plot showing differentially abundant bacterial class (Welch's t-test, Benjamini–Hochberg FDR, p<0.05) between Water_2020 and WH_2020.

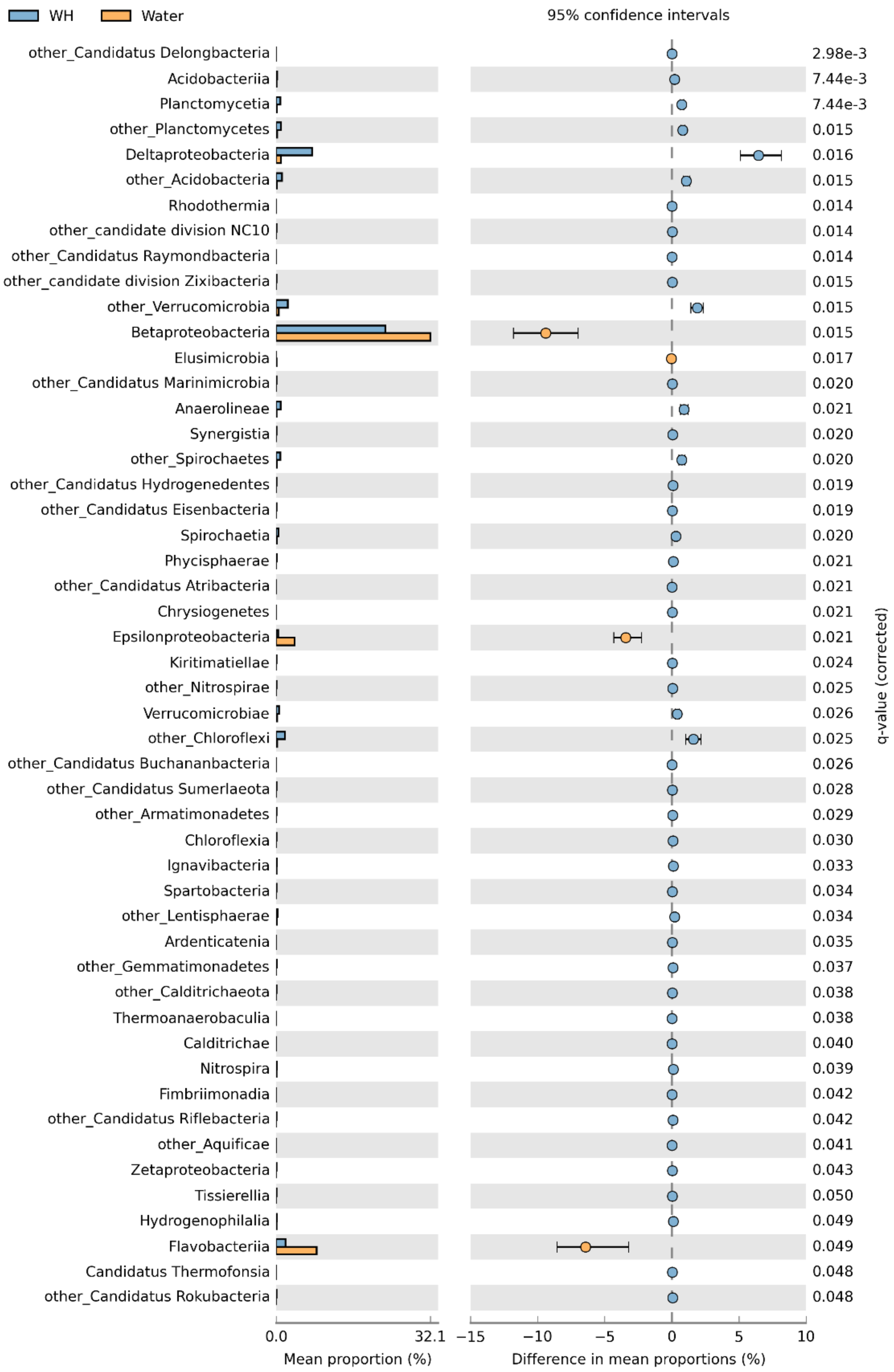


**Supplementary Figure 6.** The stacked bar illustrates top archaebacteria at class level

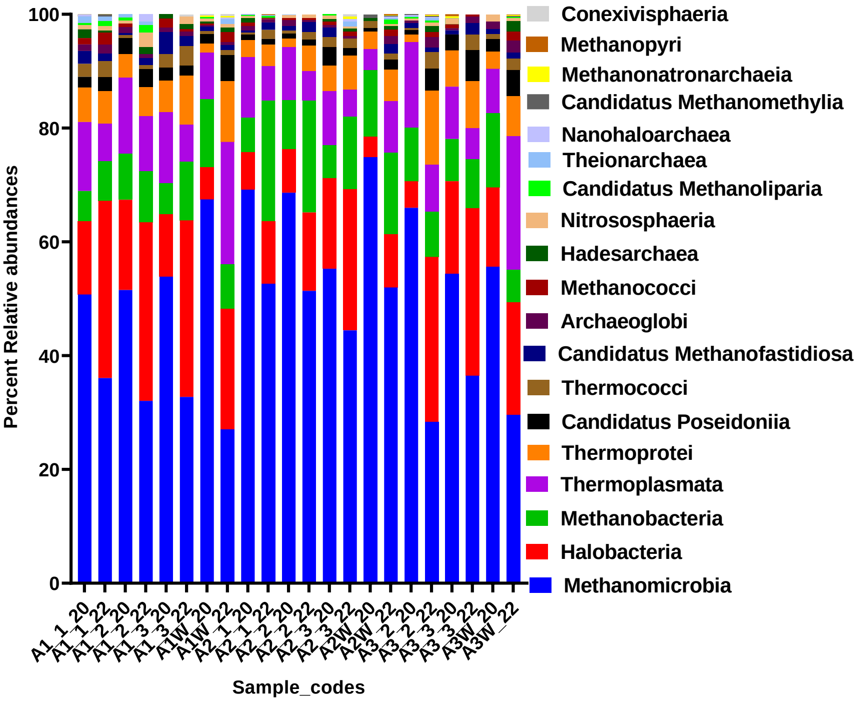


**Supplementary Figure 7A.** Box plots illustrates the alpha diversity indices chao1 and Shannon between Water_2020 and WH_2020 (t-test, p>0.05)

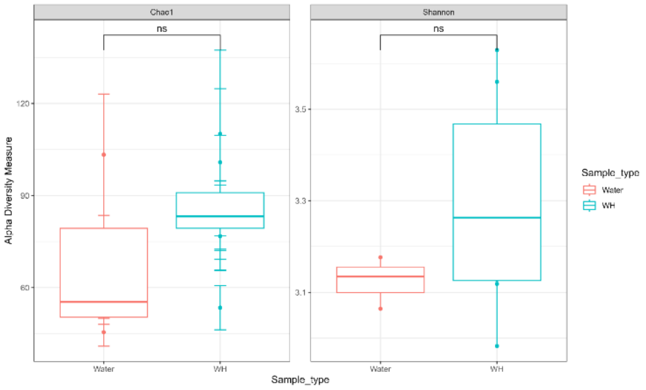


**Supplementary Figure 7B**. Box plots illustrates the alpha diversity indices chao1 and Shannon between Water_2022 and WH_2022 (t-test, p>0.05)

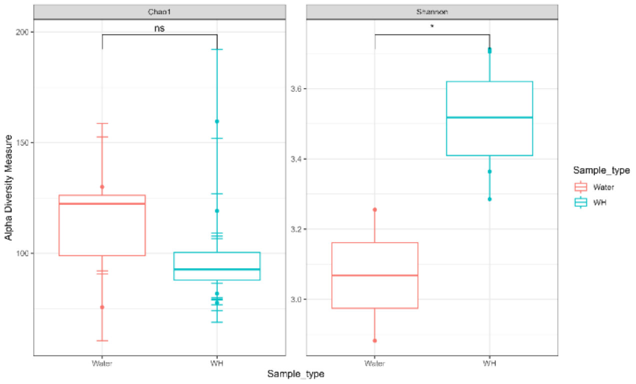


**Supplementary Figure 7C**. Box plots illustrates the alpha diversity indices chao1 and Shannon between WH_Lake and WH_2020 (t-test, p>0.05)

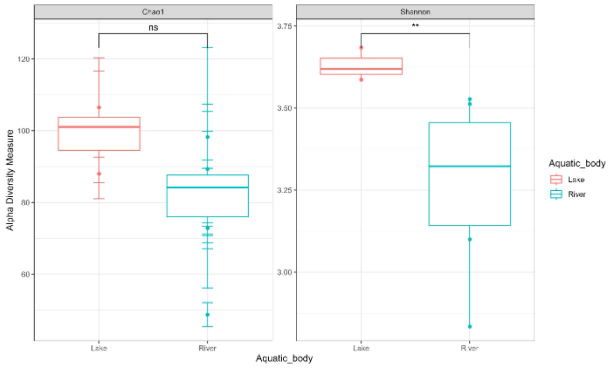


**Supplementary Figure 8A.** PCoA plots using Bray-Curtis distance between river Water_2020 and WH_2020 ($p<0.05$, R=0.53, $R^2$ =0.20, Beta-disper>0.05).

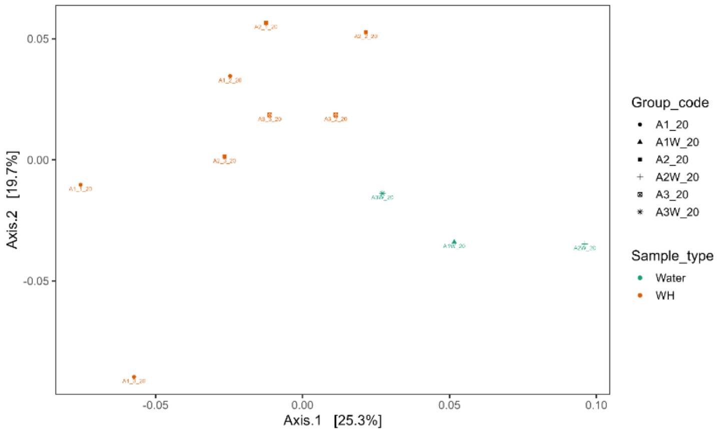


**Supplementary Figure 8B.** PCoA plots using Bray-Curtis distance between river Water_2022and WH_2022 ($p<0.05$, R=0.46, $R^2$ =0.18, Beta-disper>0.05).

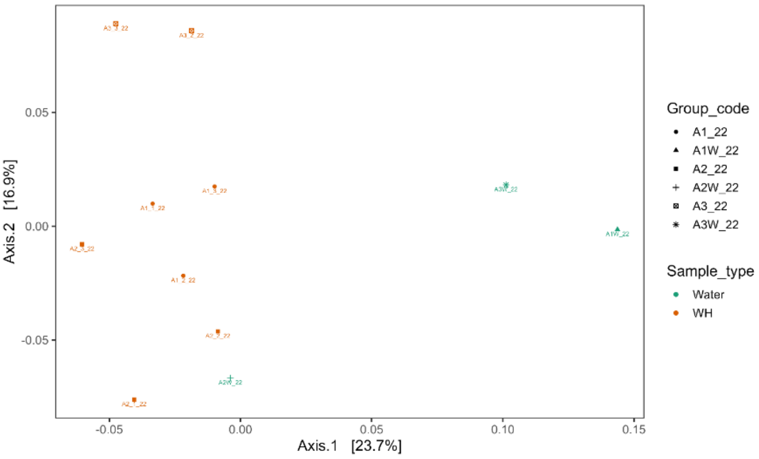


**Supplementary Figure 8C.** PCoA plots using Bray-Curtis distance between river WH_2020 and WH_Lake ($p<0.05$, R=0.94, $R^2$ =0.36, Beta-disper>0.05).

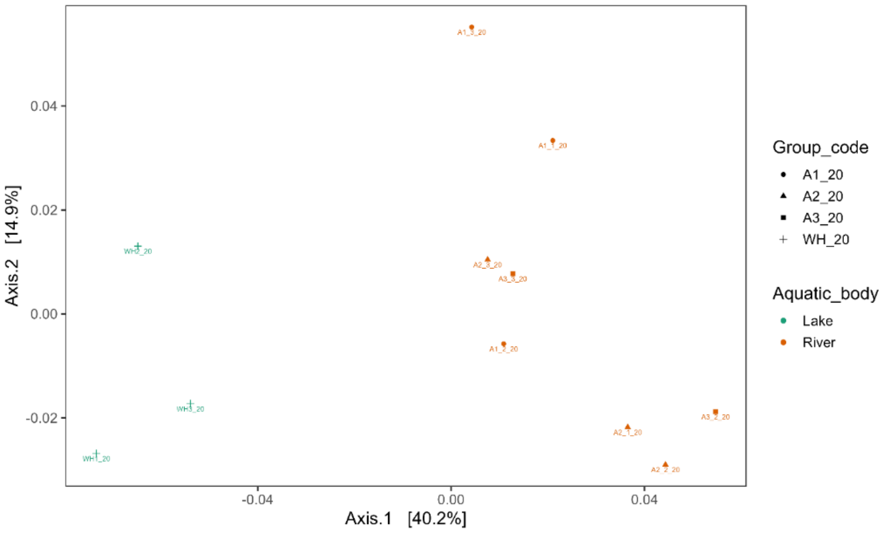


**Supplementary Figure 8D.** PCoA plots using Bray-Curtis distance between river WH_2020 and WH_2022 ($p<0.05$, R=0.42, $R^2$ =0.17, Beta-disper>0.05).

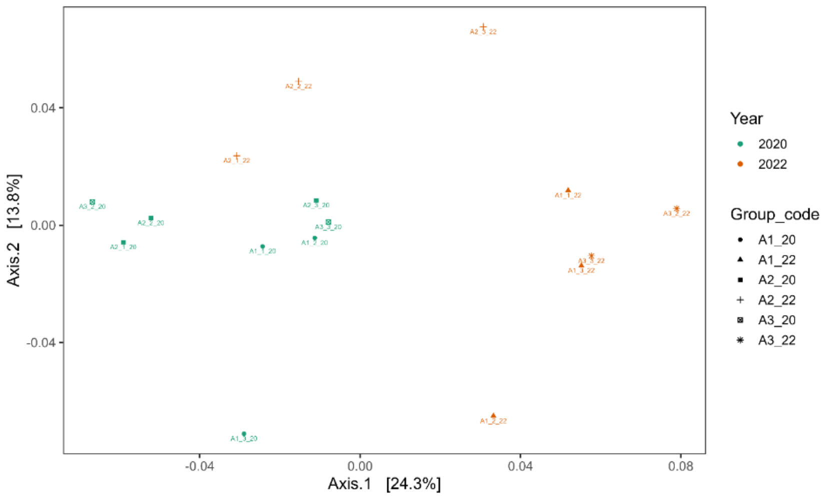


**Supplementary Figure 9.** PCoA plots using Bray-Curtis distance between river sites of WH_2022 ($p<0.05$, R=0.95, $R^2$ =0.63, Beta-disper>0.05).

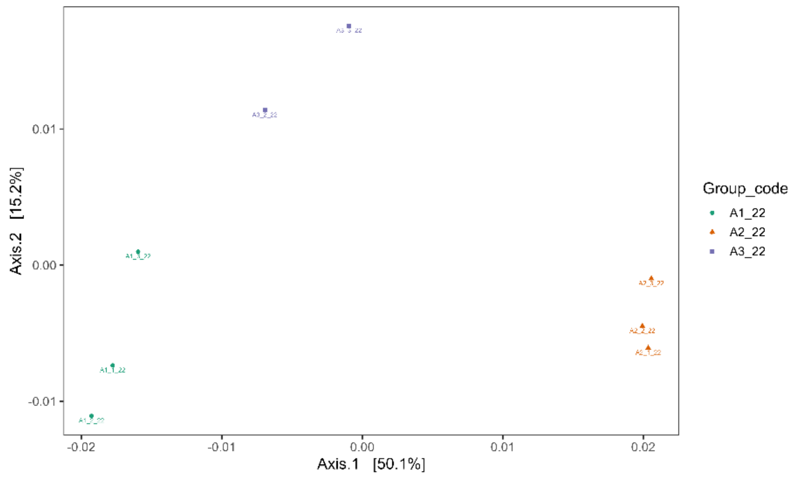


**Supplementary Figure 10.** Bacterial taxa at class level encoding for PDEs

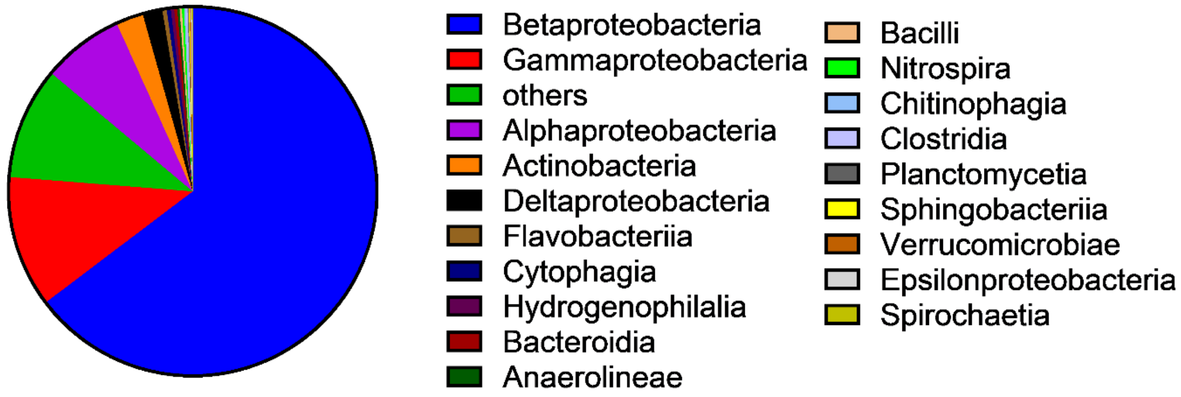


**Supplementary Figure 11.** Bacterial taxa at genus level encoding for PDEs

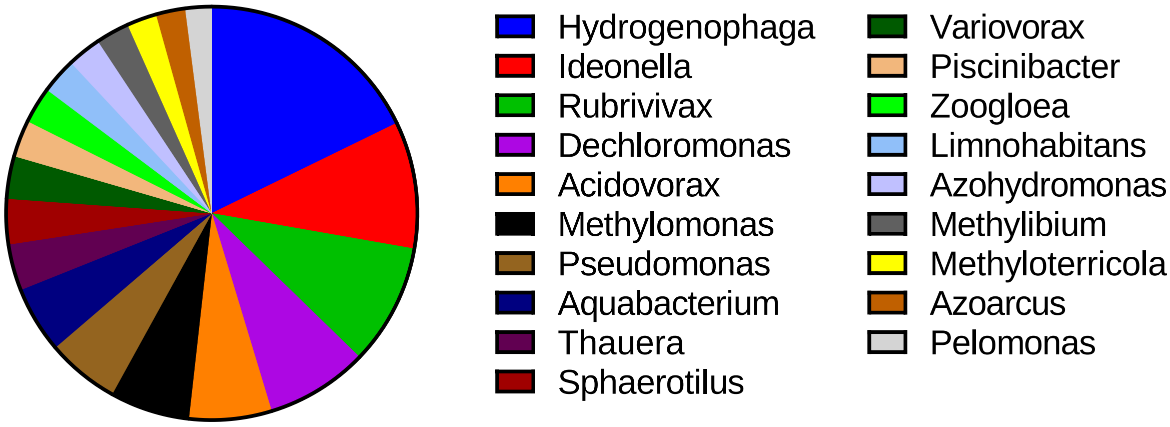


**Supplementary Figure 11**. MRGs against different biocides and metals

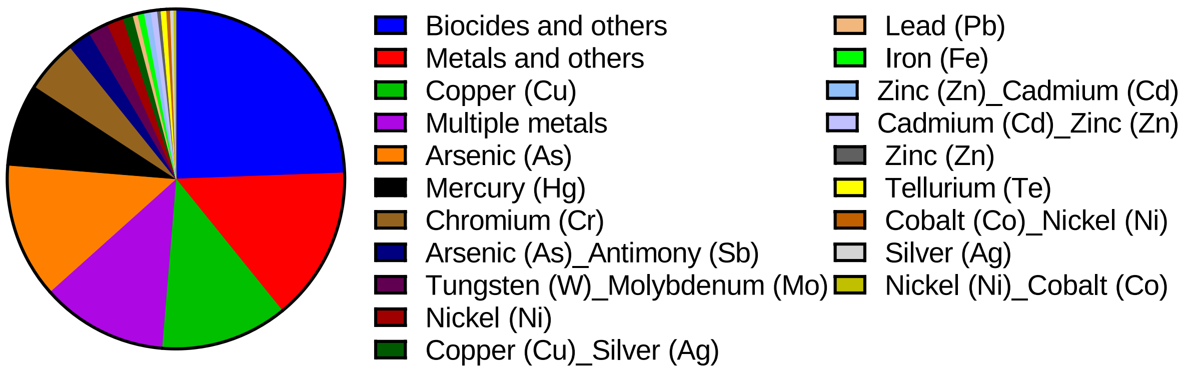


Supplementary Figure 12. Differential abundant bacterial families (White's non-paramteric test, Benjamini–Hochberg FDR, p<0.05) wh_2020 and Water_2020

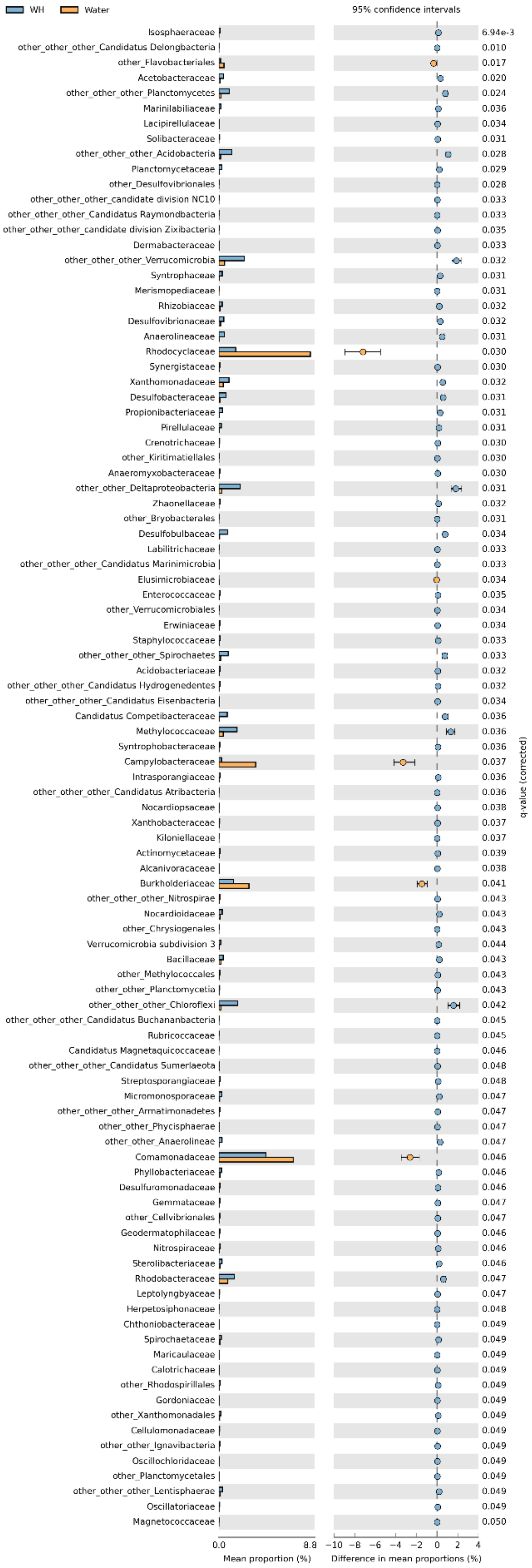

WH
Water
95% confidence intervals
Isosphaeraceae 6.94e-3
other_other_other_Candidatus Delongbacteria 0.010
other_Flavobacteriales 0.017
Acetobacteraceae 0.020
other_other_other_Planctomycetes 0.024
Marinilabiliaceae 0.036
Lacipirellulaceae 0.034
Solibacteraceae 0.031
other_other_other_Acidobacteria 0.028
Planctomycetaceae 0.029
other_Desulfovibrionales 0.028
other_other_other_candidate division NC10 0.033
other_other_other_Candidatus Raymondbacteria 0.033
other_other_other_candidate division Zixibacteria 0.035
Dermabacteraceae 0.033
other_other_other_Verrucomicrobia 0.032
Syntrophaceae 0.031
Merismopediaceae 0.031
Rhizobiaceae 0.032
Desulfovibrionaceae 0.032
Anaerolineaceae 0.031
Rhodocyclaceae 0.030
Synergistaceae 0.030
Xanthomonadaceae 0.032
Desulfobacteraceae 0.031
Propionibacteriaceae 0.031
Pirellulaceae 0.031
Crenotrichaceae 0.030
other_Kiritimatiellales 0.030
Anaeromyxobacteraceae 0.030
other_other_Deltaproteobacteria 0.031
Zhaonellaceae 0.032
other_Bryobacterales 0.031
Desulfobulbaceae 0.034
Labilitrichaceae 0.033
other_other_other_Candidatus Marinimicrobia 0.033
Elusimicrobiaceae 0.034
Enterococcaceae 0.035
other_Verrucomicrobiales 0.034
Erwiniaceae 0.034
Staphylococcaceae 0.033
other_other_other_Spirochaetes 0.033
Acidobacteriaceae 0.032
other_other_other_Candidatus Hydrogenedentes 0.032
other_other_other_Candidatus Eisenbacteria 0.034
Candidatus Competibacteraceae 0.036
Methylococcaceae 0.036
Syntrophobacteraceae 0.036
Campylobacteraceae 0.037
Intrasporangiaceae 0.036
other_other_other_Candidatus Atribacteria 0.036
Nocardiopsaceae 0.038
Xanthobacteraceae 0.037
Kiloniellaceae 0.037
Actinomycetaceae 0.039
Alcanivoracaceae 0.038
Burkholderiaceae 0.041
other_other_other_Nitrospirae 0.043
Nocardioidaceae 0.043
other_Chrysiogenales 0.043
Verrucomicrobia subdivision 3 0.044
Bacillaceae 0.043
other_Methylococcales 0.043
other_other_Planctomycetia 0.043
other_other_other_Chloroflexi 0.042
other_other_other_Candidatus Buchananbacteria 0.045
Rubricoccaceae 0.045
Candidatus Magnetaquicoccaceae 0.046
other_other_other_Candidatus Sumerlaeota 0.048
Streptosporangiaceae 0.048
Micromonosporaceae 0.047
other_other_other_Armatimonadetes 0.047
other_other_Phycisphaerae 0.047
other_other_Anaerolineae 0.047
Comamonadaceae 0.046
Phyllobacteriaceae 0.046
Desulfuromonadaceae 0.046
Gemmataceae 0.047
other_Cellvibrionales 0.047
Geodermatophilaceae 0.046
Nitrospiraceae 0.046
Sterolibacteriaceae 0.046
Rhodobacteraceae 0.047
Leptolyngbyaceae 0.047
Herpetosiphonaceae 0.048
Chthoniobacteraceae 0.049
Spirochaetaceae 0.049
Maricaulaceae 0.049
Calotrichaceae 0.049
other_Rhodospirillales 0.049
Gordoniaceae 0.049
other_Xanthomonadales 0.049
Cellulomonadaceae 0.049
other_other_Ignavibacteria 0.049
Oscillochloridaceae 0.049
other_Planctomycetales 0.049
other_other_other_Lentisphaerae 0.049
Oscillatoriaceae 0.049
Magnetococcaceae 0.050
q-value (corrected)
0.0
8.8
Mean proportion (%)
−10 −8 −6 −4 −2 0 2 4
Difference in mean proportions (%)
